# A modality gap in personal-genome prediction by sequence-to-function models

**DOI:** 10.64898/2026.02.01.702969

**Authors:** Xinming Tu, Anna Spiro, Maria Chikina, Sara Mostafavi

## Abstract

Sequence-to-function (S2F) models trained on reference genomes have achieved strong performance on regulatory prediction and variant-effect benchmarks, yet they still struggle to predict inter-individual variation in gene expression from personal genomes. We evaluated AlphaGenome on personal genome prediction in two molecular modalities—gene expression and chromatin accessibility—and observed a striking dichotomy: AlphaGenome approaches the heritability ceiling for chromatin accessibility variation, but remains far below baseline for gene-expression variation, despite improving over Borzoi. Context truncation and fine-mapped QTL analyses indicate that accessibility is governed by local regulatory grammar captured by current architectures, whereas gene-expression variation requires long-range regulatory integration that remains challenging.

## Main text

Deep learning models that map genomic DNA to regulatory readouts are widely used to interpret non-coding variation and nominate mechanisms at GWAS loci.^1–5^ These models learn transferable regulatory “grammar” from large functional genomics compendia and can generalize to unseen loci and alleles via in silico variant-effect scoring.^1–4,6–8^ However, multiple studies have shown that predicting cross-individual gene-expression variation from personal genomes remains difficult for state-of-the-art reference-trained models, with frequent failures in direction-of-effect prediction.^9,10^

AlphaGenome is a recent unified sequence model that takes ∼1 Mb of DNA sequence as input and predicts diverse regulatory modalities at high resolution, representing the current state of the art—often by a substantial margin across downstream tasks—and one of the largest deep models for learning sequence determinants of gene regulation.^11^ While AlphaGenome reports strong performance on track prediction and variant-effect benchmarks, whether such scaling translates into improved personal-genome prediction remains uncertain. Existing evaluations of sequence-to-function models largely focus on reference-genome track accuracy or single-variant in silico effect scores. While informative, these benchmarks do not directly test the core personalization objective: predicting *cross-individual molecular variation from full haplotypes* (**Fig. 1a**). Variant-centric benchmarks also often rely on high-confidence fine-mapped variant–gene links to define “gold-standard” causal variants (**Fig. 1b**), but stringent thresholds such as PIP>0.9 trade coverage for certainty; variants with moderate PIP frequently reflect LD or multi-causal architectures rather than absence of *cis*-genetic signal. Across multiple fine-mapped eQTL resources, only a minority(11–25%) of fine-mapped genes have any variant with PIP>0.9 (**Fig. 1c**). As a result, a substantial fraction of gene expression heritability remains unexplained by fine-mapped variants (**Fig. S1a-b**), motivating locus-level personal-genome evaluation approaches that avoid committing to a single causal variant and enable genome-wide assessment of predictive performance. In addition, standard variant-effect evaluations pool variant-level effects across genes and implicitly weight genes with many evaluated variants more heavily; this pooling can make overall performance appear strong while masking gene-level sign errors (i.e., incorrect direction of effect) that directly limit cross-individual expression prediction(**Fig. S1c-d**). A recent preprint evaluated AlphaGenome for personal expression prediction using GTEx and reported improved direction-of-effect prediction relative to Enformer, while noting persistent limitations.^12^ Here, we complement and extend that work by benchmarking AlphaGenome in a heritability-matched framework across two modalities—gene expression and chromatin accessibility—and by connecting modality-specific performance to regulatory distance, in a regime where model capacity, resolution, and context length have markedly increased while the underlying training data and sequence space remain largely unchanged (“reference-genome training”).

**Figure 1.**
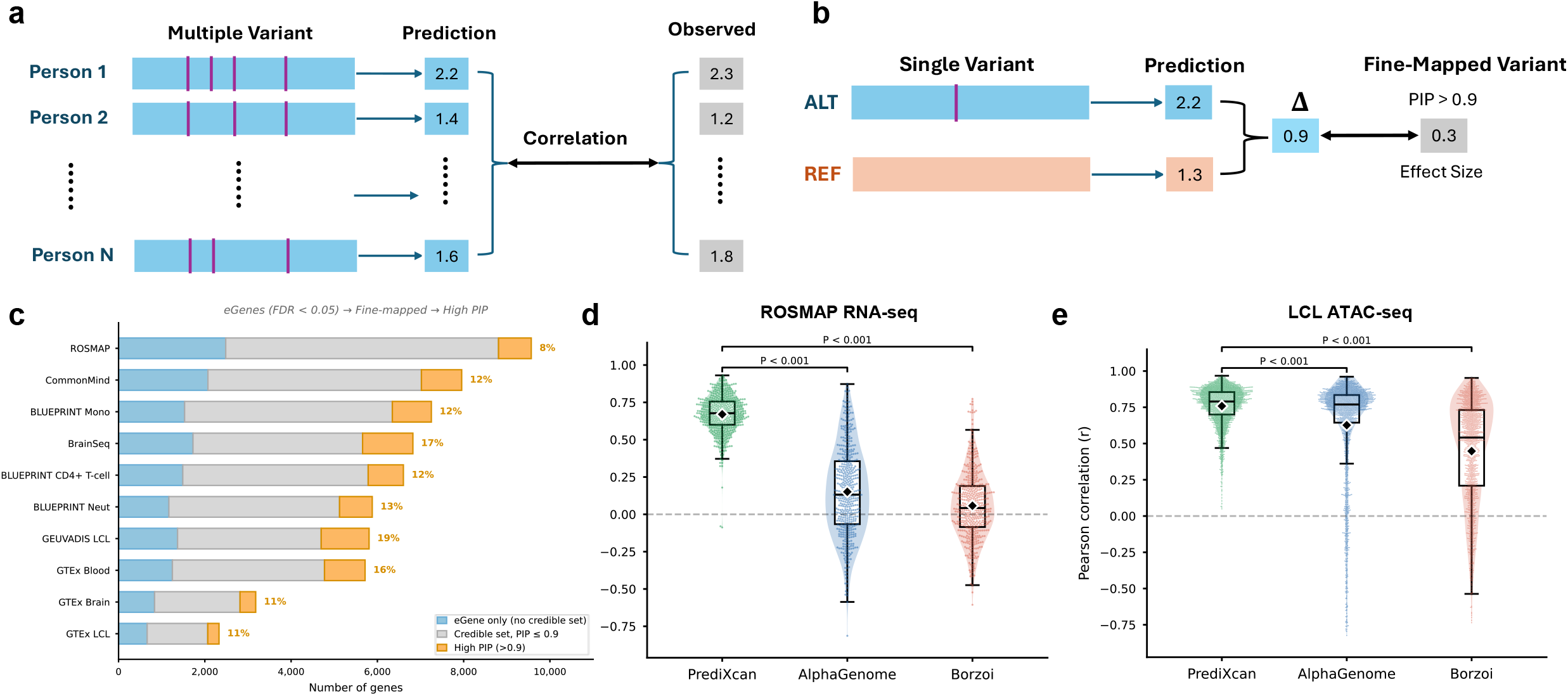
Personal genome-based benchmarking of S2F models. a) Schematic of personal-genome evaluation. For each locus, models are run on haplotype-resolved personal sequences from many individuals, each containing multiple variants within the input window. Locus-level predictions are compared to observed molecular phenotypes across individuals, and performance is summarized by cross-individual correlation (Pearson *R*). b) Schematic of standard variant-effect benchmarking. Predicted allelic effects for a single variant (difference between ALT and REF predictions, Δ) are compared to variant-level “gold-standard” effects derived from fine-mapping or population summary statistics, typically restricting evaluation to high-confidence variants (e.g., PIP>0.9). c) Limited coverage of high-confidence fine-mapped variants across studies. For multiple eQTL fine-mapping resources, bars show the number of fine-mapped genes (x-axis), partitioned into genes with at least one variant with PIP>0.9 (orange) versus genes without a variant exceeding this threshold (grey). Percentages indicate the fraction of fine-mapped genes with max PIP>0.9 in each study. d) Personal gene expression prediction across ROSMAP DLPFC genes. Distributions show per-gene Pearson correlation (r) between predicted and observed expression across held-out individuals for a cis-genotype ElasticNet baseline (PrediXcan-style), AlphaGenome, and Borzoi. e) Personal chromatin accessibility prediction across LCL ATAC peaks. Distributions show per-peak Pearson correlation (r) between predicted and observed accessibility across held-out individuals for the same methods. In d–e, each point represents one locus (peak or gene), violins show the distribution density, boxplots indicate median and interquartile range (whiskers, 1.5×IQR), and black diamonds denote the mean. P values are from paired two-sided Wilcoxon signed-rank tests comparing per-locus correlations between models.

Specifically, we evaluated personal-genome predictions (per locus and across individuals) by correlating model predictions from each individual’s diploid sequence with observed molecular measurements (*i*.*e*., in held-out individuals as the model has not seen any personal genomes prior to this assessment) in two settings: gene expression and chromatin accessibility. For gene expression, we used paired whole-genome sequencing (WGS) and dorsolateral prefrontal cortex (DLPFC) bulk RNA-seq from the ROSMAP study (n=859).^13,14^ For chromatin accessibility, we used paired WGS and ATAC-seq data from lymphoblastoid cell lines (LCLs; n=100).^15,16^ We compared AlphaGenome to Borzoi^17^ and to a *cis*-genotype elastic-net baseline (PrediXcan-style),^18^ which provides a locus-specific estimate of the linear gene expression heritability under the same cohort and preprocessing.

Because AlphaGenome inference is API-based and runtime grows steeply with context length (**Fig. S2**), we performed evaluation on targeted sets of loci enriched for *cis*-genetic signal using a held-out selection procedure (Online Methods). Across 500 genes with high genetic heritability in DLPFC (ROSMAP), AlphaGenome achieved significantly higher cross-individual correlations than Borzoi but remained far below the *cis*-genotype baseline (**Fig. 1d**). Mean per-gene Pearson *R* was 0.151 for AlphaGenome, 0.058 for Borzoi, and 0.670 for the genotype baseline (all pairwise Wilcoxon signed-rank p-value<0.001; **Fig. 1d**). Gene-wise performance was only weakly concordant between AlphaGenome and the genotype-based heritability baseline (*R*=0.17; p-value<0.001; **Fig. S3a**) whereas concordance was higher between AlphaGenome and Borzoi (*R*=0.34; p-value<0.001), suggesting that current deep learning models share inductive biases and predictive structure yet fail to capture much of the *cis*-genetic component of expression that a gene-specific linear model learns.

AlphaGenome provides multiple RNA output tracks relevant to brain tissues, including ENCODE-annotated and GTEx-annotated tracks.^19,20^ Repeating the evaluation across these track families yielded consistent results (mean *R*=0.139 versus max *R*=0.151; per-gene correspondence *R*=0.67; **Fig. S4a-c**), indicating that poor personal-expression performance is not explained by a simple track-choice artifact.

In contrast, AlphaGenome performed substantially better on personal chromatin accessibility prediction. Across 1,852 heritable LCL ATAC peaks, AlphaGenome reached mean *R*=0.627, closer to the genotype-based heritability baseline *R*=0.758 and outperformed Borzoi (mean *R*=0.448; all pairwise P<0.001; **Fig. 1e**). As in expression, performance was more concordant between the two deep models (AlphaGenome–Borzoi *R*=0.59) than between AlphaGenome and the genotype baseline (*R*=0.15; **Fig. S3b**), consistent with shared deep-model inductive biases.

We also assessed whether the accessibility result depended on track selection. AlphaGenome provides 44 donor-specific LCL ATAC tracks, and peak-level accuracy was highly stable across these tracks (mean r≈0.62; range ≈0.01 across tracks;**Fig. S4d**). In contrast, Borzoi showed substantially greater sensitivity to track choice across candidate LCL chromatin tracks (best mean r≈0.44; range ≈0.22; **Fig. S4e**); nevertheless, Borzoi remained below AlphaGenome even under favorable track selection.

Together, these results reveal a clear modality gap: scaling and architectural advances deliver substantially improved personal-genome prediction for chromatin accessibility but not for gene expression. We hypothesized that this gap reflects how causal regulatory signals are distributed across genomic distances. Chromatin accessibility measured by ATAC-seq is closely tied to local transcription factor binding and nucleosome positioning,^16^ whereas gene expression integrates promoter activity with distal enhancer regulation mediated by 3D genome organization and enhancer–promoter coupling.^19,21^ To probe distance dependence, we performed context truncation experiments by restricting the effective sequence window provided to each model. While gene expression prediction improved with increasing context but remained far below the genotype baseline across all contexts tested (**Fig. 2a**), accessibility predictions saturated rapidly: performance remained high even at short 4KB contexts (**Fig. 2b**). Notably, longer context improved cross-region (across-genomic-position) agreement between predicted and observed profiles for both modalities (**Fig S5a-b**), suggesting that models leverage long context to recover broad spatial patterns while still failing to predict cross-individual expression variation. At the same time, both deep models capture between-locus differences in mean activity (mean predicted vs. mean observed across genes/peaks), even when cross-individual correlations are low (**Fig. S6**)

**Fig. 2.**
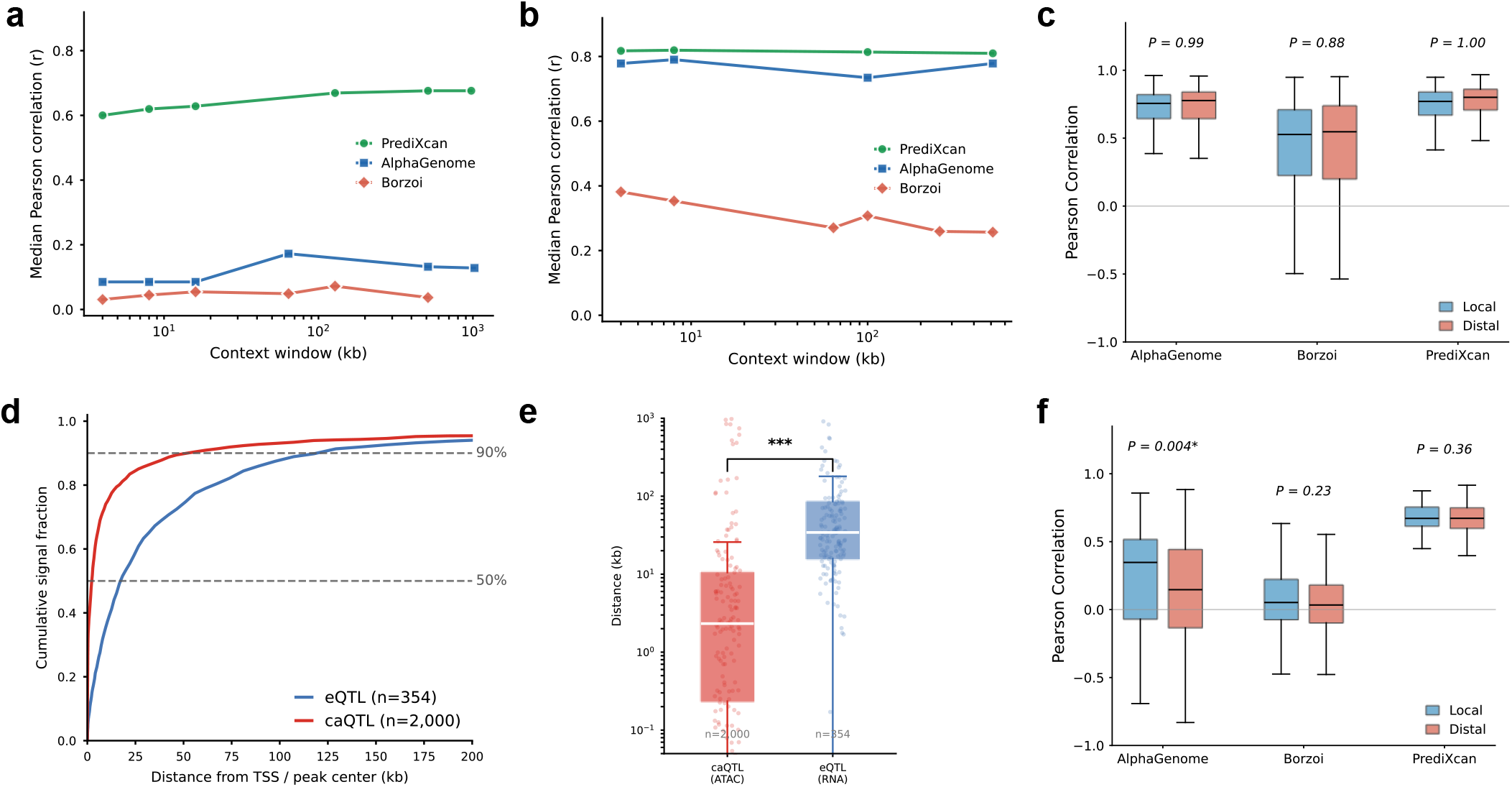
Regulatory distance shows modality-specific context dependence and local–distal stratification. a) Gene expression: median cross-individual prediction accuracy (Pearson r; median across loci) as a function of sequence context window length. Points indicate evaluated context lengths (4 kb–1 Mb); lines connect medians across the same gene set with valid predictions at all contexts (n = 462 genes). b) Chromatin accessibility: median cross-individual prediction accuracy as a function of context window length for LCL ATAC peaks with valid predictions across all contexts (n = 1,988 peaks). c) Accessibility prediction stratified by regulatory localization. Peaks are split into “local” versus “distal” groups using caQTL lead-variant distance to peak center (median split; Methods). Boxplots show per-peak cross-individual accuracy for each method; P values above each method compare local vs distal groups (Mann–Whitney U test; Methods). d) Cumulative localization of QTL signal as a function of distance. For eQTLs (blue), curves summarize cumulative fine-mapped posterior mass around transcription start sites (TSSs) across genes; for caQTLs (red), curves summarize cumulative localization around peak centers across peaks. Horizontal dashed lines indicate 50% and 90% cumulative signal. e) Distribution of regulatory-distance metrics for caQTLs and eQTLs. caQTL distance is the absolute lead-variant distance to peak center (n = 2,000 peaks); eQTL distance is d90, the minimal distance from the TSS capturing 90% of cumulative fine-mapped posterior mass (n = 354 genes). P value compares distance distributions (Wilcoxon rank-sum test; Methods). f) Expression prediction stratified by regulatory localization. Genes are split into “local” versus “distal” groups using d90 (local: d90 < 20 kb; distal: d90 ≥ 20 kb; Methods). Boxplots show per-gene cross-individual accuracy for each method; P values compare local vs distal groups (Mann–Whitney U test; Methods).

To connect these observations to genetic architecture, we quantified the spatial distribution of causal QTL signals using fine-mapped QTL resources. For caQTLs we considered lead-variant distances to peak centers^15^, and for eQTLs we summarized the distance distribution of fine-mapped posterior mass around transcription start sites (TSSs) using a d90 statistic (distance capturing 90% of cumulative posterior mass). caQTL signal was highly local (median lead distance 2.3 kb), whereas eQTL signal was substantially more distal (median d90 34 kb; **Fig. 2d-e**).

Consistent with a long-range integration limitation, AlphaGenome personal-expression prediction was significantly worse for distally regulated genes than for locally regulated genes (P=0.004; **Fig. 2f**) as defined by fine-mapped variants, while chromatin accessibility prediction showed no detectable local-versus-distal difference (P≈1 across methods; **Fig. 2c**). These results suggest that current S2F models learn highly predictive local regulatory grammar, sufficient for personal accessibility, but do not learn the enhancer–gene wiring required to map distal regulatory elements to target genes with the fidelity needed to predict cross-individual expression.

Finally, we asked *why* zero-shot personal-expression prediction is so poor. One possibility is that the pretrained sequence model fails to encode genotype-dependent information relevant to expression. An alternative is that genotype-dependent signal *is* present in the model-derived features, but the pretrained RNA output head does not combine these features into accurate gene-level expression differences across individuals.Training supervised locus-specific readouts on model-derived features substantially increased held-out performance; strikingly, even features from an untrained (random-weight) Borzoi network supported non-trivial prediction (**Fig. S7**). This diagnostic is consistent with substantial genotype-related information being preserved under high-dimensional featurizations, while the end-to-end pretrained mapping remains misaligned for zero-shot cross-individual expression prediction.

In summary, across two personal-genome benchmarks we find that AlphaGenome substantially improves over Borzoi yet remains far below heritability-matched genotype baselines for gene expression, while approaching those baselines more closely for chromatin accessibility. These findings complement prior reports that reference-trained deep models underperform on personal transcriptome variation^9,10^ and indicate that simply scaling context length is insufficient to solve the gene-expression problem. We hypothesize that closing this gap likely requires progress along two complementary axes. First, models may need stronger inductive biases and supervision for enhancer–promoter wiring, for example using objectives informed by 3D genome organization and perturbation-validated enhancer–gene maps.^19,21^ Second, reference-genome training alone may be fundamentally limited for personal-genome generalization; incorporating population-scale sequence diversity via haplotype-resolved genomes or pangenome references^22^ and/or QTL-informed objectives may be necessary to teach models how real allele combinations modulate regulatory circuitry across individuals.

## Acknowledgments

We are grateful to Žiga Avsec and Anshul Kundaje for their thoughtful comments and valuable feedback on this manuscript.

## Online Method

### Data and code availability

ROSMAP genotype and dorsolateral prefrontal cortex (DLPFC) bulk RNA-seq data were accessed under controlled access via Synapse (accession to be provided) consistent with ROSMAP data use requirements. LCL ATAC-seq peak quantifications and caQTL fine-mapping results were obtained from Kumasaka et al. Phased LCL genotypes were obtained from the 1000 Genomes Project Phase 3 resource. Fine-mapped eQTL credible sets (variant-level posterior inclusion probabilities, PIPs) were obtained from the eQTL Catalogue (accession QTD000434 for ROSMAP DLPFC eQTLs; additional accessions listed below for the fine-mapping coverage analysis) and, when not available there, from original study repositories.

All analysis code—including preprocessing pipelines, personal-genome sequence construction, model inference wrappers, statistical analyses, and figure-generation scripts—is available in our public repository: **github.com/mostafavilabuw/alphagenome-personal-eval**. This resource will be maintained and finalized upon publication

### Study design

We benchmarked S2F models for zero-shot personal-genome prediction across two molecular modalities with distinct regulatory architectures: bulk RNA-seq gene expression (ROSMAP DLPFC) and ATAC-seq chromatin accessibility (LCLs). For each locus (gene or ATAC peak), we evaluated cross-individual prediction accuracy as the Pearson correlation between predicted and observed molecular phenotypes across individuals in a held-out test set. We compared AlphaGenome and Borzoi to a cohort-matched *cis*-genotype ElasticNet baseline (PrediXcan-style). No statistical methods were used to predetermine sample size; sample sizes were determined by data availability. Unless otherwise stated, analyses used a fixed random seed (42).

## Datasets and preprocessing

### ROSMAP DLPFC RNA-seq and genotypes

We analyzed paired genotype and bulk RNA-seq data from the Religious Orders Study and Memory and Aging Project (ROSMAP). The dataset comprised n = 859 individuals with matched phased genotypes (GRCh38/hg38 coordinates) and DLPFC bulk RNA-seq.

Observed expression phenotypes were gene-level, covariate-adjusted log-TPM values. Briefly, expression was summarized at the gene level, converted to TPM, log-transformed, and residualized using linear regression to remove known covariates. Covariates included age at death, sex, post-mortem interval, RNA integrity number (RIN), technical batch factors, and the top genotype principal components (10 PCs unless otherwise noted). Residualized values were used as the observed expression phenotype for all predictive models and statistical baselines.

### LCL ATAC-seq, caQTL annotations, and genotypes

We used processed ATAC-seq chromatin accessibility measurements from lymphoblastoid cell lines (LCLs) derived from the 1000 Genomes Project Phase 3 British cohort (GBR; n = 100 individuals), as distributed by Kumasaka et al.15 Accessibility was provided as GC-corrected, log-FPKM–normalized peak values with 16 principal components regressed out to mitigate technical variation. Peak coordinates were defined on GRCh37/hg19. Unless otherwise stated, analyses were restricted to autosomal peaks. Matched phased genotypes for the same 100 individuals were obtained from the 1000 Genomes Project Phase 3 resource.

### Genotype processing and quality control

For cohort-specific genotype matrices used in statistical baselines, we retained bi-allelic SNPs passing standard quality control: per-variant call rate > 0.95, per-individual call rate > 0.95, and minor allele frequency (MAF) thresholds described below (ROSMAP: MAF ≥ 0.01; LCL: MAF ≥ 0.05 for ElasticNet training). For personal-genome sequence construction used in deep-model inference, phased small variants (SNPs and short indels) were incorporated (see below). Structural variants and records with symbolic alternate alleles were excluded. Copy number variants were not modeled.

### Train/validation/test splits and locus selection

To focus evaluation on loci with detectable *cis*-genetic regulation while avoiding selection bias toward any deep model, we used held-out selection based on the *cis*-genotype baseline.

### ROSMAP splits and gene selection

ROSMAP individuals (n = 859) were split into mutually exclusive sets: training (n = 689), validation (n = 85), and test (n = 85). We trained *cis*-genotype ElasticNet models (PrediXcan-style; described below) for genes in the training set and ranked genes by the Pearson correlation between predicted and observed expression in the validation set. The top 500 genes by validation performance were carried forward for model benchmarking. All reported comparisons between AlphaGenome, Borzoi, and the genotype baseline for gene expression were computed exclusively in the held-out test set (n = 85 individuals) using these 500 genes.

### LCL splits and ATAC peak selection

LCL individuals (n = 100) were split into training (n = 80) and test (n = 20). Within the training set, we performed 5-fold cross-validation of *cis*-genotype ElasticNet models to rank peaks by mean cross-validated Pearson correlation. Peaks with strongest *cis*-genetic predictability were prioritized for benchmarking. We report results on the held-out test set (n = 20) for peaks with complete data and valid predictions across methods; the primary benchmark set contained 1,852 peaks.

### Personal genome sequence construction

Reference genome sequence was extracted from GRCh38/hg38 for ROSMAP analyses and from GRCh37/hg19 for LCL analyses.

For each individual and each locus, we constructed two haplotype-specific DNA sequences (maternal and paternal) by applying phased genotypes to the reference sequence within the input window centered at the relevant anchor point (gene transcription start site (TSS) for expression; ATAC peak center for accessibility). Variants were applied in genomic order. Variants whose reference allele did not match the reference genome sequence at the stated coordinate were excluded.

Because insertions and deletions alter coordinate mappings between reference and haplotype sequences, we tracked cumulative coordinate offsets for each haplotype to maintain correct mapping between reference coordinates and haplotype sequence coordinates. This mapping was used to aggregate predictions over gene bodies and peak intervals after indel-induced shifts.

Sequences shorter than the required input length were center-padded with ‘N’ bases; longer sequences were center-cropped to the model-specific input length. Predictions were generated separately for each haplotype and averaged to produce a diploid prediction per individual.

## Prediction models and inference

### AlphaGenome inference (API-based)

AlphaGenome predictions were obtained via the AlphaGenome API. For each locus and individual, we supplied haplotype-resolved DNA sequence centered on the anchor point (TSS for genes; peak center for ATAC). Unless otherwise noted, the default context window was 100 kb. For context-dependence analyses, we evaluated total input lengths of 4 kb, 16 kb, 100 kb, 500 kb, and 1 Mb (see below).

### Gene expression (RNA tracks)

For gene expression, we used AlphaGenome RNA-seq tracks annotated as relevant to brain/DLPFC, including ENCODE-annotated forward and reverse strand tracks and a GTEx-annotated track. For ENCODE, we summed forward and reverse predictions to obtain a strand-combined signal. For each track, we aggregated base- or bin-level predictions over the annotated gene body interval (TSS to transcription end site) by summation and applied a log1p transformation. Main-text results report the GTEx-annotated track unless stated; ENCODE-based results were analyzed in parallel (Supplementary).

### Chromatin accessibility (DNASE tracks)

For chromatin accessibility, we used AlphaGenome chromatin-accessibility output (DNASE) tracks annotated as LCL. AlphaGenome provides 44 donor-specific LCL accessibility tracks; for each peak we summed predictions over the full ATAC peak interval and applied log1p. We computed cross-individual Pearson correlation separately for each track and summarized peak-level accuracy by the mean across tracks. Track-by-track variability is reported in Supplementary analyses.

### Borzoi inference (Flashzoi implementation)

We evaluated Borzoi using a FlashAttention-optimized implementation (“Flashzoi”) that preserves identical model outputs while improving inference efficiency. Borzoi takes a fixed 524,288-bp DNA sequence as input and outputs predictions in 32-bp bins across the central 196,608 bp. Where applicable, we averaged predictions across the four model replicates provided with Borzoi.

Input sequences were one-hot encoded; ambiguous bases (‘N’) were encoded as a uniform distribution across A/C/G/T. For each locus and individual, maternal and paternal haplotype sequences were generated, Borzoi inference was run separately on each haplotype, and outputs were averaged to produce a diploid prediction.

For RNA-seq predictions, we selected brain-relevant RNA tracks based on Borzoi track metadata and summed strand-specific predictions where required before aggregation. For chromatin predictions, we evaluated LCL-relevant chromatin tracks (DNase/ATAC where available) and performed track-sensitivity analyses (Supplementary). Locus-level predictions were obtained by summing binned predictions overlapping each gene body or ATAC peak interval, followed by log1p transformation.

### *Cis*-genotype ElasticNet baseline (PrediXcan-style)

As a heritability-matched statistical baseline, we trained *cis*-genotype ElasticNet regression models in each cohort to predict residualized molecular phenotypes from *cis* genetic variation. For each gene/peak, we included bi-allelic SNP genotypes within ±100 kb of the anchor point (TSS for genes; peak center for ATAC peaks). Genotypes were coded as allele dosages (0/1/2). ElasticNet models were trained using scikit-learn with l1_ratio = 0.5 and max_iter = 2000. Regularization strength (alpha) was selected from a log-spaced grid spanning 0.001 to 10 (20 values). For ROSMAP, hyperparameters were selected using the validation set; for LCL, hyperparameters were selected using 5-fold cross-validation within the training set. Only variants passing cohort-specific MAF thresholds were included (ROSMAP: MAF ≥ 0.01; LCL: MAF ≥ 0.05). Baseline predictions were evaluated only in held-out test individuals.

### Context dependence (context truncation experiments)

To assess dependence on regulatory distance, we repeated personal-genome evaluation across multiple total sequence context lengths: 4 kb, 16 kb, 100 kb, 500 kb, and 1 Mb, each centered on the locus anchor point (gene TSS for expression; ATAC peak center for accessibility). For AlphaGenome, we requested the corresponding input length via the API.

For Borzoi, which requires a fixed 524,288-bp input sequence, we emulated shorter effective contexts (4 kb–500 kb) by embedding the extracted sequence at the center of the required input and padding the flanks with ‘N’ bases; the native Borzoi input served as the maximum context (reported as ∼500 kb for comparability). Context analyses were performed on the subset of loci with valid predictions across all evaluated contexts for each method.

## Fine-mapped QTL localization and regulatory distance metrics

### eQTL fine-mapping and d90

To quantify the spatial extent of causal regulation for gene expression, we used fine-mapped eQTL credible sets from the eQTL Catalogue (accession QTD000434), which provides SuSiE-based posterior inclusion probabilities (PIPs) for putatively causal variants. For each gene with an available credible set, we computed d90: the minimal absolute distance from the TSS required to capture 90% of the cumulative PIP mass when variants are sorted by increasing |distance to TSS|. For summary curves, we computed cumulative PIP as a function of distance for each gene and reported the mean across genes.

### caQTL localization

To quantify causal localization for chromatin accessibility, we used caQTL fine-mapping outputs from the LCL ATAC-seq study. For peaks with high-confidence lead variants (PLead ≥ 0.5), we computed lead-variant distance as the absolute distance between the lead variant position and the ATAC peak center (midpoint of peak start and end). For cumulative distance curves, we reported the empirical cumulative distribution of lead-variant distances across peaks.

### Local-versus-distal stratification

For genes, we defined local regulation as d90 < 20 kb and distal regulation as d90 ≥ 20 kb. For peaks, we stratified by lead-variant distance using a median split (threshold = 326 bp in this study). Stratified performance comparisons were conducted using test-set cross-individual prediction accuracy for each locus.

### Fine-mapping coverage under high-confidence (PIP > 0.9) filtering

To quantify how restricting evaluation to high-confidence fine-mapped variants impacts benchmark coverage, we summarized gene-level fine-mapping resolution across multiple independent *cis*-eQTL fine-mapping resources (Fig. 1c). We analyzed fine-mapped credible sets from ten studies spanning brain, whole blood, and immune cell types: GTEx v8 brain frontal cortex BA9 (QTD000176), GTEx v8 whole blood (QTD000356), GTEx v8 lymphoblastoid cell lines (QTD000221), ROSMAP dorsolateral prefrontal cortex (QTD000434), GEUVADIS lymphoblastoid cell lines, BLUEPRINT monocytes, BLUEPRINT neutrophils, BLUEPRINT CD4+ T cells, CommonMind Consortium dorsolateral prefrontal cortex, and BrainSeq dorsolateral prefrontal cortex. Fine-mapping outputs (credible sets with variant-level PIPs) were obtained from the eQTL Catalogue when available and otherwise from original study repositories.

For each study, we defined the set of fine-mapped genes as genes with at least one reported fine-mapped variant (i.e., appearing in any credible set with a non-missing PIP). Many genes have multiple independent *cis*-eQTL signals and thus multiple credible sets; to summarize fine-mapping resolution per gene, we computed maxPIP, defined as the maximum PIP across all variants from all credible sets for that gene within the study. We then reported the number and fraction of genes with maxPIP > 0.9.

### Cross-dataset variant concordance analysis

To evaluate the consistency of AlphaGenome variant-effect scoring across independent fine-mapped cis-eQTL resources, we performed a cross-dataset concordance analysis using fine-mapped credible sets from ten studies spanning brain, blood, and immune cell types (from the eQTL Catalogue when available and otherwise from original study repositories). For each study, we restricted to genes with exactly one high-confidence variant (defined as having exactly one variant with posterior inclusion probability (PIP) > 0.9 across all credible sets for that gene within that study). For each retained variant–gene pair, we extracted the observed fine-mapped effect size (β) from the fine-mapping output and obtained an AlphaGenome signed variant score for the same variant using a 1 Mb sequence context centered on the gene transcription start site (TSS). We quantified agreement between observed and predicted effects using (i) Pearson correlation between β and AlphaGenome scores within each study, and (ii) direction concordance, defined as the fraction of variant–gene pairs where the sign of the observed effect matched the sign of the AlphaGenome score: concordant = (sign(β) = sign(score)), concordance rate = mean(concordant).

These analyses were performed independently for each study and summarized across datasets

### Top causal variant analysis

To assess how much cis-genotype predictive signal is captured by a single variant and to quantify the coverage–performance tradeoff imposed by restricting to high-confidence fine-mapped variants, we compared three variant-based prediction strategies for ROSMAP DLPFC gene expression within the same train/validation/test split framework described above. All strategies used cis variants within ±100 kb of the gene TSS, with genotypes encoded as allele dosages (0/1/2), and performance was evaluated per gene as test-set cross-individual Pearson correlation between predicted and observed residualized expression.

- Full cis-genotype ElasticNet (PrediXcan-style): the primary baseline using all eligible cis variants within ±100 kb and ElasticNet regularization/hyperparameter selection as described above.
- Top-beta single-variant model: for each gene, we identified the single cis variant with the largest absolute coefficient in the fitted full ElasticNet model and generated predictions using only that variant (i.e., the corresponding single-variant linear predictor derived from the full model fit). This analysis quantifies how much of the full multi-variant baseline is captured by the strongest single variant selected by regularized regression.
- High-PIP restricted model: for each gene, we restricted the cis variant set to variants with PIP > 0.9 from SuSiE fine-mapping (eQTL Catalogue credible sets) and evaluated prediction using this restricted variant set. Because many genes have no cis variant exceeding PIP > 0.9, genes lacking any high-PIP variant were treated as having no available high-confidence predictor under this restriction and were assigned a null performance value (r = 0) for summary visualizations; performance summaries conditioned on high-PIP availability were computed on the subset of genes with at least one PIP > 0.9 variant (Supplementary).

### Supervised readouts on model-derived representations

To test whether heritable signal is present in model-derived representations but not expressed by pretrained output heads, we trained supervised locus-specific readouts using features derived from model outputs. For each gene, we constructed feature vectors from model-predicted chromatin signal summarized in fixed-width bins (256 bp) across the input window, applied log1p, and standardized features using training-set statistics. We trained an ElasticNet readout to predict observed residualized expression in the training set and evaluated held-out test performance. As a negative control, we repeated the procedure using features derived from an untrained (random-weight) Borzoi network. These analyses are reported in Supplementary Fig. S6.

## Evaluation metrics and statistical analysis

### Cross-individual accuracy

For each gene or peak, cross-individual prediction accuracy was defined as the Pearson correlation coefficient (r) between predicted and observed phenotypes across individuals in the held-out test set. Correlations were computed using scipy.stats.pearsonr. We summarized locus-wise performance using distributions across genes/peaks, including mean and standard deviation (Fig. 1).

### Method comparisons and concordance

To compare methods on the same set of loci, we used paired two-sided Wilcoxon signed-rank tests over per-locus correlation values (for example, AlphaGenome vs Borzoi across the same genes). To quantify concordance of locus-wise performance between methods, we computed Pearson correlation between per-locus accuracies across genes/peaks.

### Distance-distribution and stratified comparisons

To compare regulatory distance distributions between caQTL and eQTL measures, we used a two-sided Wilcoxon rank-sum test. For local-versus-distal stratified performance comparisons, we used one-sided Mann–Whitney U tests with the alternative hypothesis that locally regulated loci have higher prediction accuracy than distally regulated loci.

### Runtime benchmarking

Where indicated (Supplementary), wall-clock inference time per sequence was measured over repeated runs for AlphaGenome (end-to-end API runtime including overhead) and for Borzoi local inference (Flashzoi; fixed 524,288-bp input). Reported values are means across repeated measurements with run-to-run variability summarized by standard deviation.

---

**Supplementary Fig. 1.**
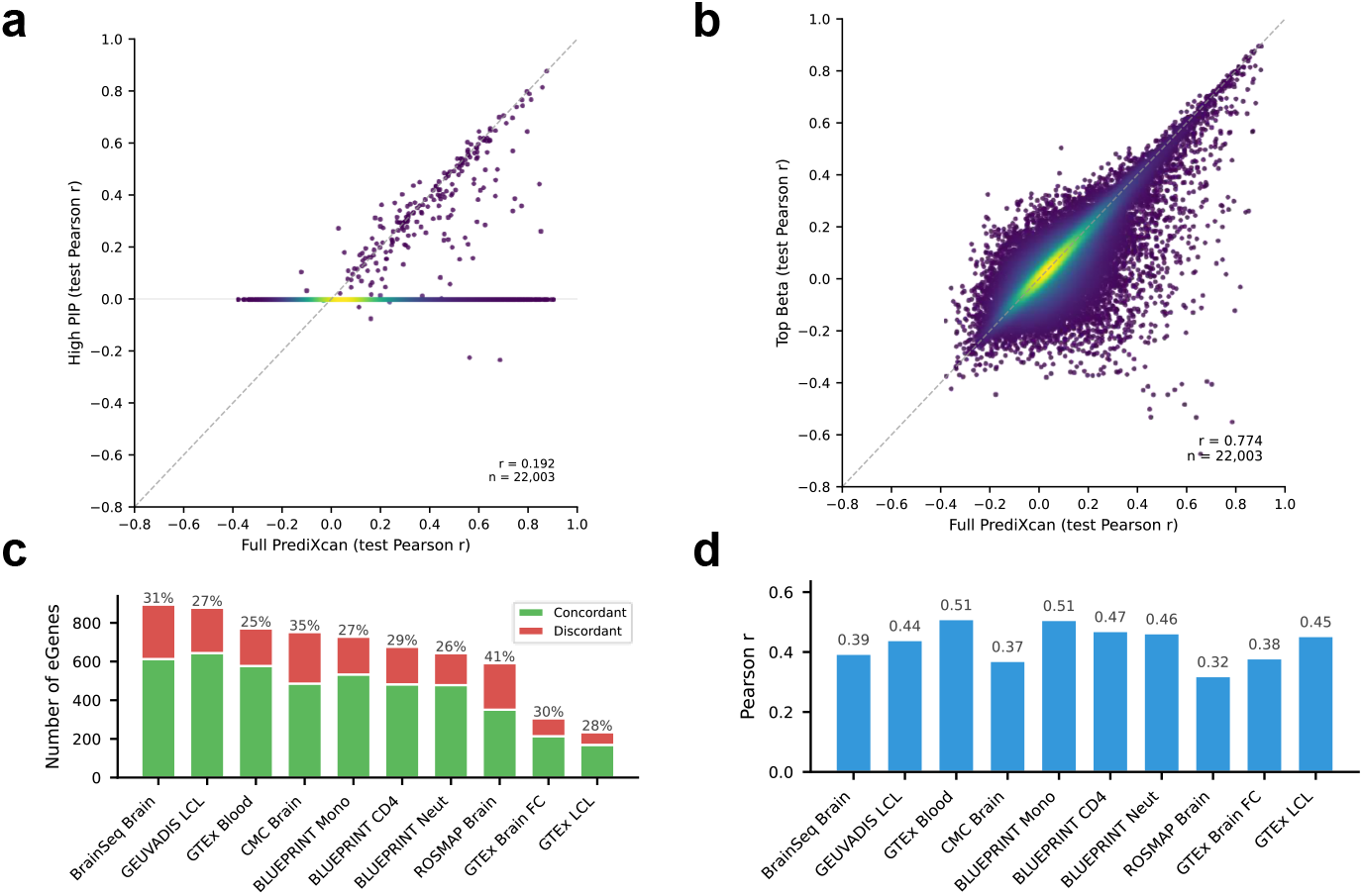
Fine-mapping stringency limits cis signal capture and pooled variant-effect metrics can mask gene-level direction errors. **a**, Gene-wise agreement between a single-variant predictor restricted to a high-confidence fine-napped variant (“High PIP”; e.g., PIP > 0.9; Methods) and the full cis-genotype ElasticNet baseline (“Full PrediXcan”), quantified as test-set cross-individual Pearson correlation (r) for each gene (n = 22,003). Many genes lack a qualifying high-PIP variant and therefore contribute near-zero predictive accuracy under this restriction. **b**, Agreement for the same comparison using a single-variant predictor based on the strongest cis variant (“Top $\beta$”; Methods). Points are colored by density; dashed line indicates y = x. **c**, Gene-level direction-of-effect concordance across fine-mapped cis-eQTL resources. For genes with a nominated high-confidence causal variant (Methods), stacked bars show the number of eGenes with concordant (green) versus discordant (red) effect direction between the model-predicted allelic effect and the reported eQTL effect; percentages denote the discordant fraction in each study. **d**, Pooled variant-effect size agreement for the same nominated causal variants. Bars show the Pearson correlation between predicted and observed signed effect sizes (Δ prediction versus eQTL effect estimate; Methods) across variants within each resource; values above bars report the corresponding correlations.

**Supplementary Fig. 2.**
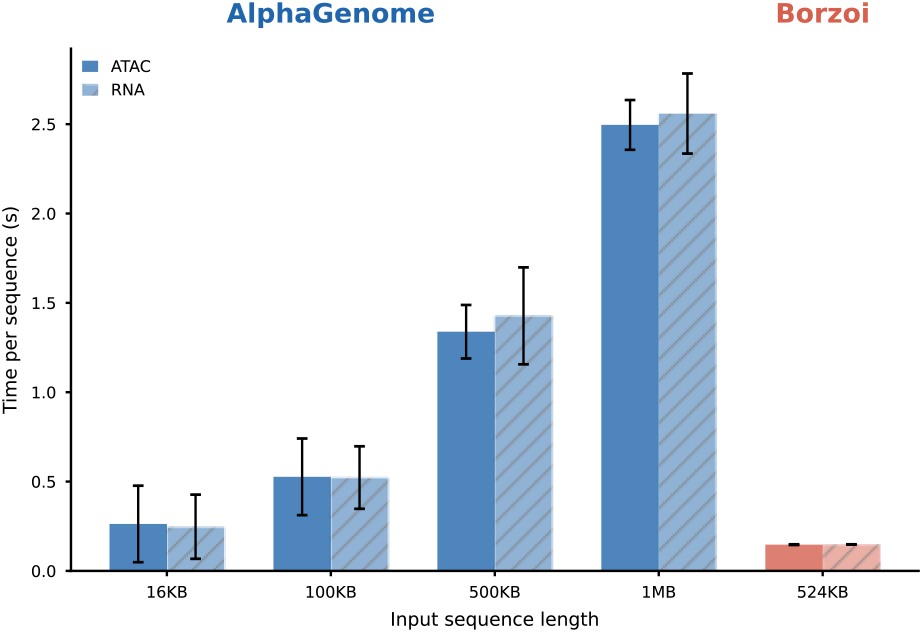
Inference time scales steeply with context for AlphaGenome. Wall-clock inference time per sequence as a function of input context length for AlphaGenome (API, including network transmission overhead; RNA setting for ROSMAP and ATAC setting for LCL) and for Borzoi (local, single A4000 GPU; fixed 524 kb input). Bars indicate means; error bars indicate s.d. across runs.

**Supplementary Fig. 3.**
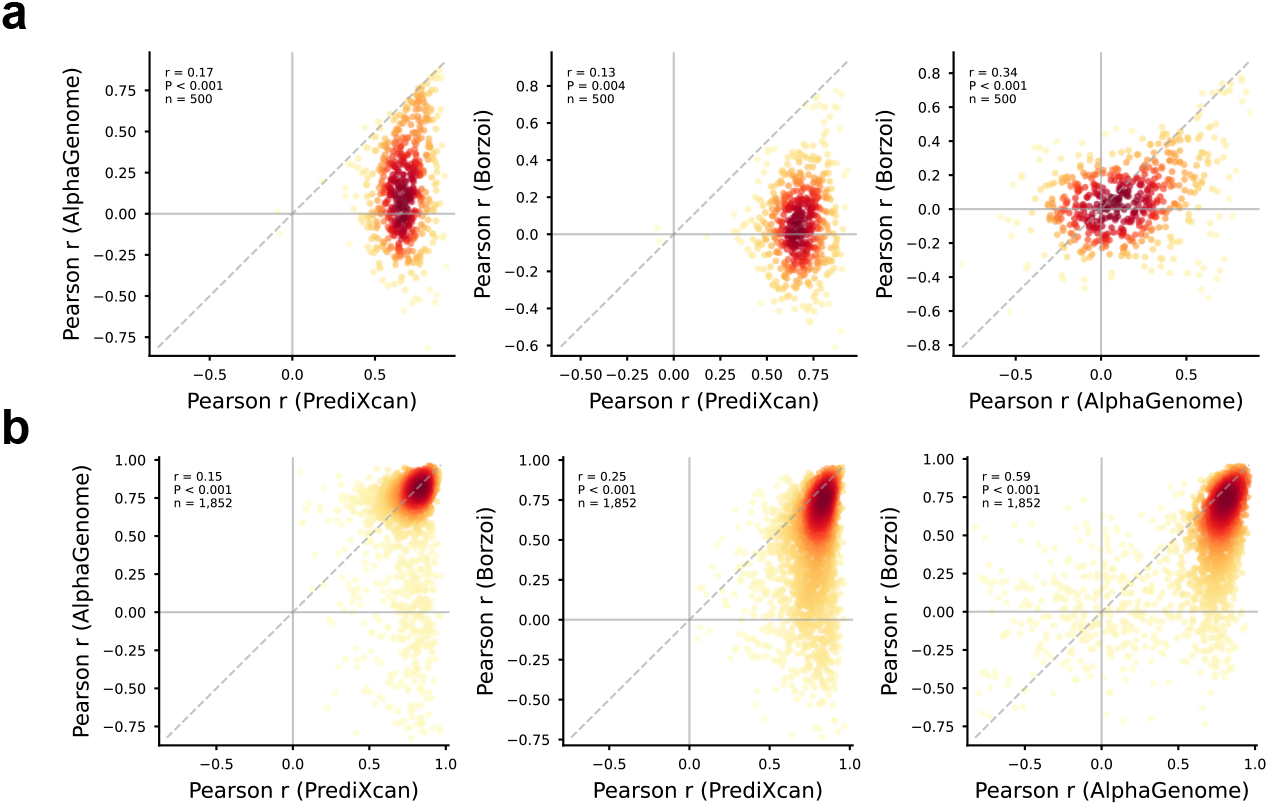
Pairwise comparison of per-gene accuracies across methods. **a)** Pairwise comparison of per-gene accuracies across methods for the same 500 genes. Each point denotes one gene; dashed line indicates equality. Insets report Pearson correlation between per-gene accuracies (method–method concordance). b) Pairwise comparison of per-peak accuracies across methods for the same 1,852 peaks; dashed line indicates equality; insets report Pearson correlation between per-peak accuracies.

**Supplementary Fig. 4.**
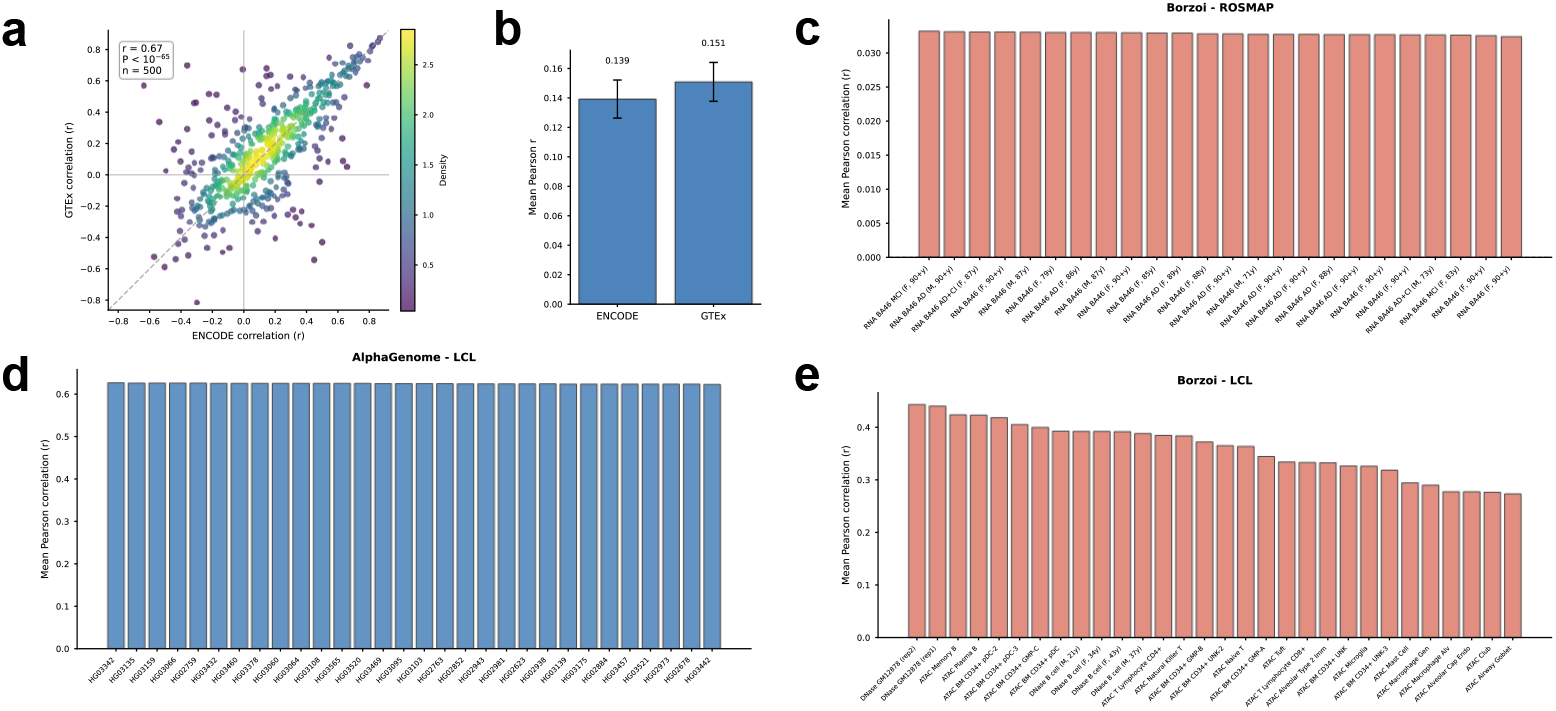
Track-choice sensitivity analyses for RNA and ATAC outputs. a) Concordance of AlphaGenome per-gene cross-individual accuracy when using two alternative DLPFC-relevant RNA-seq track families (GTEx-annotated versus ENCODE-annotated). Each point denotes one gene (n = 500); axes show per-gene Pearson r across individuals in the test set; inset reports correlation between per-gene accuracies across track families. b) Mean cross-individual accuracy for AlphaGenome under the GTEx-annotated versus ENCODE-annotated RNA track choices; bars show mean ± s.e.m. across genes (n = 500). c) Borzoi ROSMAP RNA track sweep. Each bar denotes one Borzoi RNA track considered “brain-relevant”; height is mean per-gene cross-individual accuracy across genes (n = 500), illustrating limited sensitivity to track choice under uniformly low performance. d) AlphaGenome LCL ATAC track sweep. Each bar denotes one donor-specific LCL ATAC track (44 tracks total); height is mean per-peak cross-individual accuracy across peaks (n = 1,852), showing stability across track choices. e) Borzoi LCL chromatin track sweep. Each bar denotes one candidate LCL chromatin track; height is mean per-peak cross-individual accuracy across peaks (n = 1,852), showing greater track heterogeneity than AlphaGenome.

**Supplementary Fig. 5.**
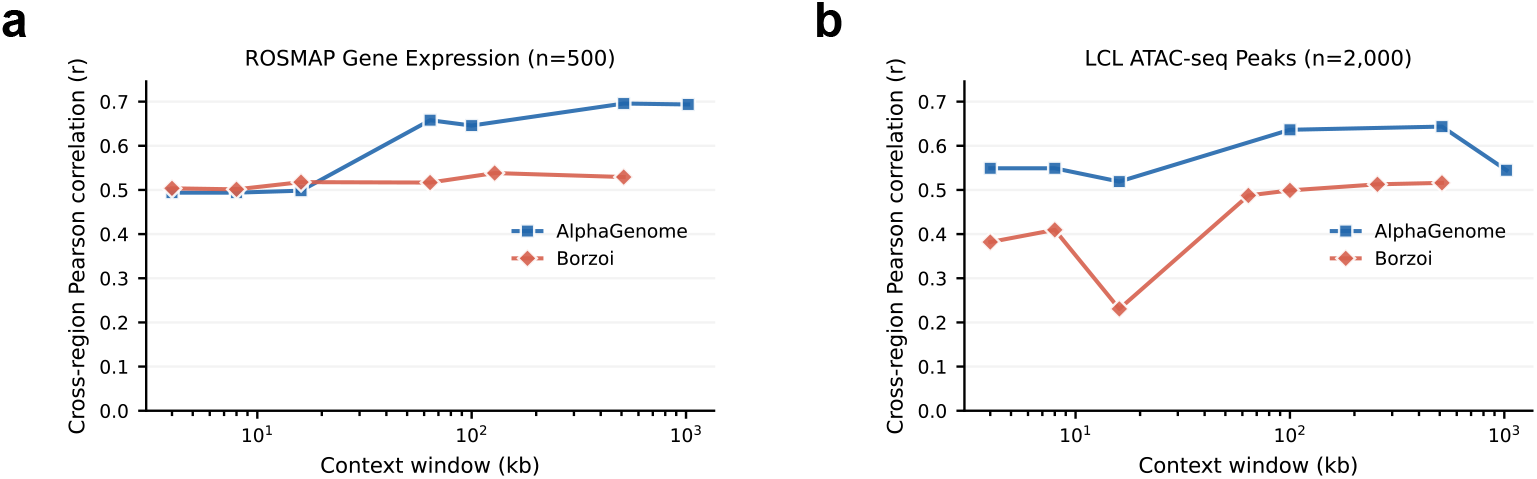
Longer context improves predicted–observed agreement across genomic positions within loci. a) ROSMAP gene-expression profiles (n = 500 genes). Cross-region agreement is computed as the Pearson correlation between predicted and observed profiles across genomic bins/positions within each gene locus, summarized across genes, as a function of context window length. **b)** LCL ATAC-seq profiles (n = 2,000 peaks). Cross-region agreement between predicted and observed profiles across genomic bins/positions within each peak, summarized across peaks, as a function of context window length. Curves are shown for AlphaGenome and Borzoi.

**Supplementary Fig. 6.**
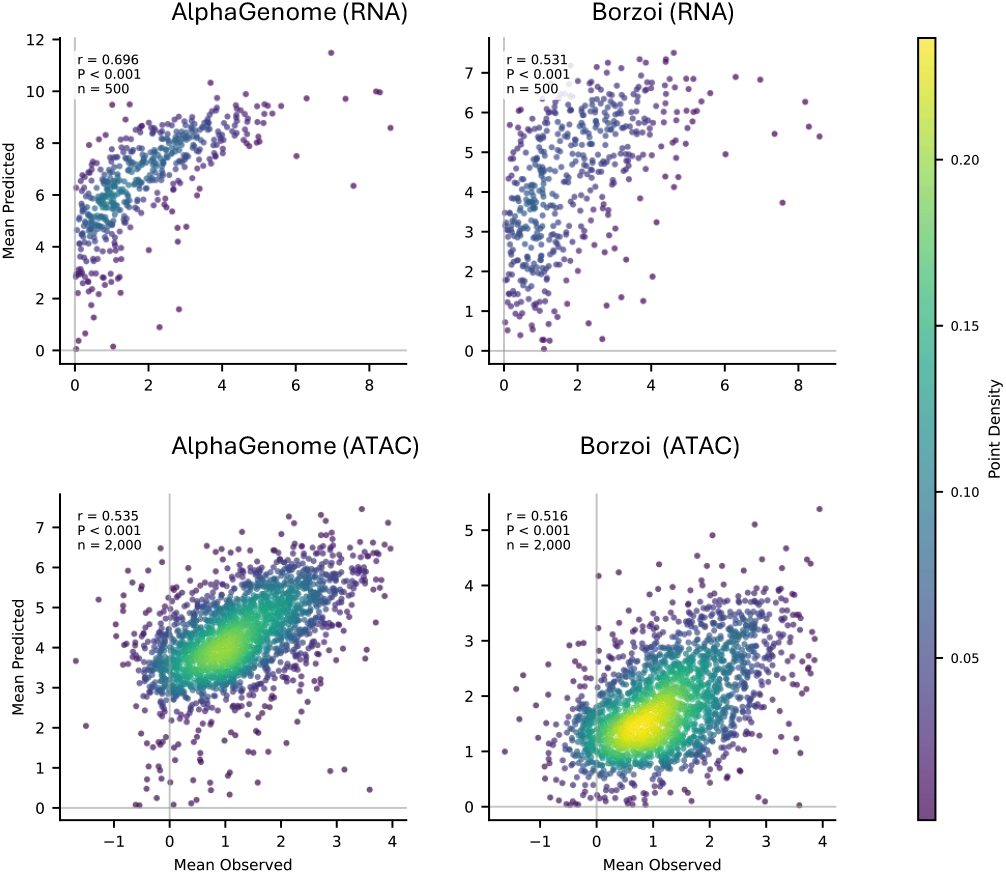
Deep models capture between-locus mean signal across genes/peaks. Scatter plots comparing mean predicted versus mean observed signal for each locus, computed across individuals in the test set. Each point denotes one gene (ROSMAP; n = 500) or one ATAC peak (LCL; n = 2,000). Panels show best track of AlphaGenome and Borzoi for each modality; insets report Pearson correlation across loci. Point color indicates local point density.

**Supplementary Fig. 7.**
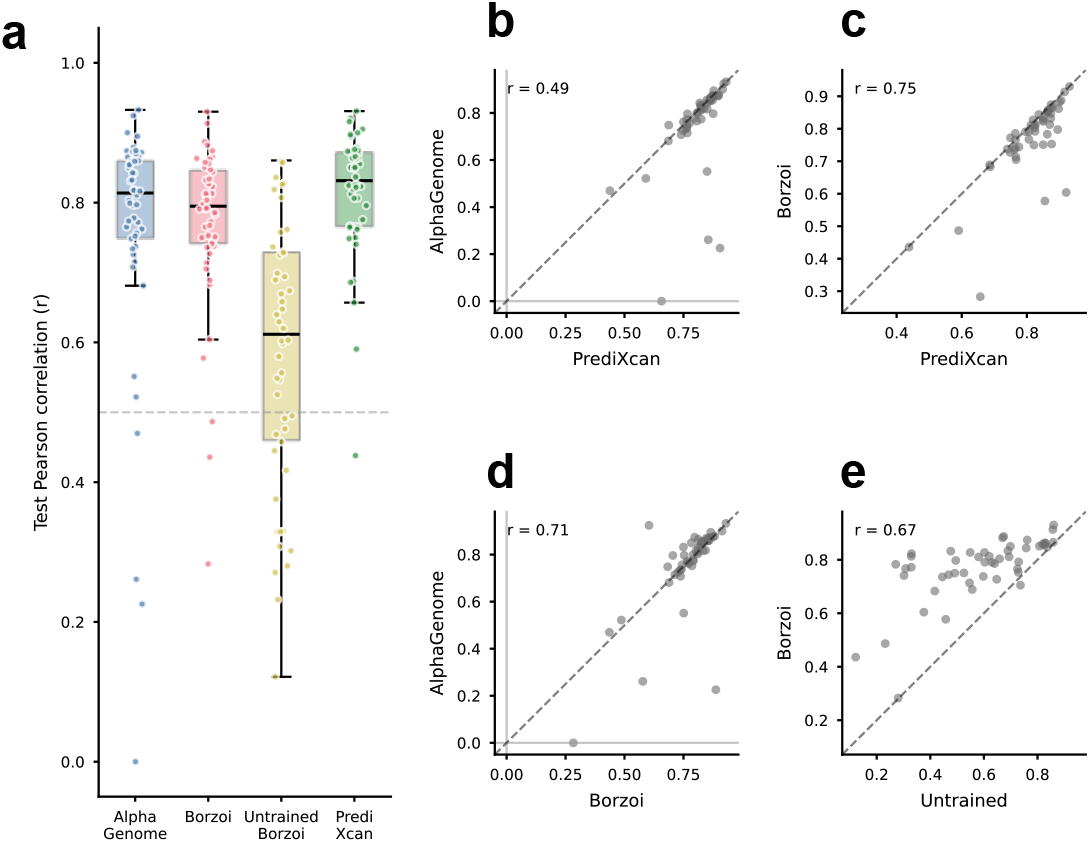
Supervised locus-specific readouts recover predictability from model-derived representations. **a)** Held-out test performance of supervised locus-specific readouts trained on model-derived features, compared to end-to-end zero-shot predictions and a cis-genotype baseline. For each gene, an elastic-net readout is trained on features derived from AlphaGenome, Borzoi, or an untrained (random-weight) Borzoi network (Methods). Boxplots show the distribution of per-gene Pearson r across individuals in the test set; points denote individual genes. **b-e)** Pairwise comparisons of per-gene test accuracies for the supervised readouts and genotype baseline. Each point denotes one gene; dashed line indicates equality. Insets report Pearson correlation across genes between the per-gene accuracies.

